# Dietary Bovine Milk Exosomes Elicit Changes in Microbial Communities in C57BL/6 Mice

**DOI:** 10.1101/356048

**Authors:** Fang Zhou, Henry A. Paz, Jiang Shu, Mahrou Sadri, Juan Cui, Samodha C. Fernando, Janos Zempleni

**Author notes:** Address correspondence to Janos Zempleni. Present address: Department of Animal and Dairy Sciences, Mississippi State University, Starkville, Mississippi, USA.

## Abstract

Exosomes and exosome-like vesicles participate in cell-to-cell communication in animals, plant and bacteria. Dietary exosomes in bovine milk are bioavailable in non-bovine species, but a fraction of milk exosomes reaches the large intestine. We hypothesized that milk exosomes alter the composition of the gut microbiome in mice. C57BL/6 mice were fed AIN-93G diets, defined by their content of bovine milk exosomes and RNA cargos: exosome/RNA depleted (ERD) versus exosome/RNA-sufficient (ERS) diets. Feeding was initiated at age three weeks and cecum content was collected at ages 7, 15 and 47 weeks. Microbial communities were identified by 16S *rRNA* gene sequencing. The dietary intake of exosomes and age had significant effects on the microbial communities in the cecum. At the phylum level, the abundance of *Verrucomicrobia* was greater in mice fed ERD compared to ERS, and the abundance of both *Firmicutes* and *Tenericutes* was smaller in mice fed ERD compared to ERS at age 47 weeks. At the family level, the abundance of *Anaeroplasmataceae* was greater in mice fed ERD compared to ERS, and the abundance of *Bifidobacteriaceae*, *Lachnospiraceae*, and *Dehalobacteriaceae* was significantly greater in mice fed ERS than mice fed ERD at age 15 weeks. Exosome feeding significantly altered the abundance of 52 operational taxonomic units; diet effects were particularly strong in the *Lachnospiraceae*, *Ruminococcaceae* and the *Verrucomicrobiaceae* families. We conclude that exosomes in bovine milk alter microbial communities in non-bovine species, suggesting that exosomes and their cargos participate in the crosstalk between bacterial and animal kingdoms.

**IMPORTANCE:** Virtually all living cells, including bacteria communicate through exosomes, which can be found in all body fluids. Exosomes and the RNA cargos have been implicated in all aspects of health and disease, *e.g.,* metastasis of cancer, neuronal signaling and embryonic development. Previously, we reported that exosomes and their microRNA cargos are not solely derived from endogenous synthesis, but may also be obtained from dietary sources such as bovine milk in non-bovine mammals. Here, we report for the first time that bovine milk exosomes communicate with the intestinal microbiome and alters microbial communities in mice. This is the first report suggesting that the gut microbiome facilitates the signaling by dietary exosomes across kingdoms: animal (cow) → bacteria → animal (mouse).

---

Exosomes are nanoparticles that play essential roles in cell-to-cell communication (1, 2). Exosomes transfer diverse cargos from donor cells to adjacent or distant recipient cells. Cargos include various species of RNA, proteins, and lipids. In recipient cells, exosome cargos alter gene expression and metabolism. For example, miR-30d secreted by the endometrium is taken up by the pre-implantation embryo and modifies the transcriptome in human embryos (3). Recently, we made the paradigm-shifting discovery that exosomes and their RNA cargos are not exclusively derived from endogenous synthesis, but can also we obtained from dietary sources. We demonstrated that human and rat intestinal cells transport exosomes from bovine milk by endocytosis and secrete microRNA cargos across the basolateral membrane (4). Similar transport mechanism operates in human vascular endothelial cells (5). Exosomes accumulate in immune cells if transferred across species boundaries (6–8). We further demonstrated that bovine milk exosomes contribute to the body pool of microRNAs in human milk feeding and murine exosome depletion studies (6, 9, 10). MicroRNAs in dietary exosomes alter gene expression across species boundaries (6, 11). These discoveries are consistent with observations by other investigators. For example, Mathivanan and co-workers and Ochiya and co-workers presented evidence in support of the bioavailability of milk exosomes at a conference on Dietary Extracellular Vesicles hosted by the International Society of Extracellular Vesicles in January of 2017 at La Trobe University, Australia. Ongoing phenotyping studies in our laboratory suggest that dietary depletion of bovine milk exosomes causes a loss in fecundity and spatial learning and memory in mice, alters plasma cytokine profiles in humans and mice, and causes an aberrant flux of purine metabolites in mice (12–14).

Despite the compelling evidence in support of the theory that milk exosomes are bioavailable, concerns have been raised that the amount of cargos, particularly microRNAs, delivered by bovine exosomes to host organisms is exceedingly low in transgenic mouse models ((15, 16), reviewed in (17)). In a recent opinion paper we suggested that, while studies of dietary microRNAs are important and warrant investigation, the controversy surrounding the field of dietary microRNAs must not impede the rate of discovery in areas such as dietary exosomes, exosome cargos other than microRNAs and non-canonical RNA signaling pathways (18). Here we tested the hypothesis that bovine milk exosomes alter microbial communities in the murine cecum and any such changes might contribute to changes in the murine hepatic transcriptome. This hypothesis was based on the following rationale. First, prokaryotic and eukaryotic microbes communicate with their environment through exosome-like vesicles (19). This observation includes gram-positive bacteria, which use vesicles for communication despite the cell wall posing a barrier for vesicle transport (20, 21). Viruses may participate in exosome signaling through hijacking and modifying exosomes (22). Second, up to 20% and 40% of RNA sequence reads in plasma from healthy adults map to bacterial and fungal genomes, respectively (23). Third, evidence suggests that orally administered, fluorophore-labeled exosomes from bovine milk are delivered to peripheral tissues (24). While that study lacked important controls (unlabeled exosomes, free fluorophore), its findings are largely consistent with our ongoing studies suggesting that endogenously and exogenously labeled milk exosomes accumulate in liver and spleen, but that a considerable fraction of orally administered exosomes escapes absorption and reaches the large intestine (25).

## RESULTS

### Microbial communities

Combinatorial effects of independent variables on microbial communities were statistically significant for the diet *x* age *x* sex interaction (*P*=0.039; Table 1). At the phylum level, the abundance of *Verrucomicrobia* was greater in mice fed an AIN-93G-based, Exosome and RNA-depleted (ERD) diet compared to Exosome and RNA-sufficient (ERS) diet (*P*=0.030). Also, *Firmicutes* were significantly more abundant in female mice fed the ERS diet compared with females fed the ERD diet at age 47 weeks (*P*<0.05). Some of the changes at the phylum level involved diet *x* age interactions. For example, *Firmicutes* and *Tenericutes* were significantly less abundant in mice fed the ERD diet compared with ERS at age 47 weeks, but not at ages 7 and 15 weeks. The abundance of *Actinobacteria* was greater (*P*=0.041) in mice fed the ERS diet compared with ERD at age 15 weeks. No significant changes at the phylum level were detected when sex was used as the sole independent variable. All statistical analyses are shown in Table S1.

**TABLE 1.** PERMANOVA analysis of the effect of diet, age and sex on microbial communities in the cecum of mice.

|  | **Df** | **SumsOfSqs** | **MeanSqs** | **F. Model** | **R<sup>2</sup>** | **P value** |
| --- | --- | --- | --- | --- | --- | --- |
| **Diet x age x sex** | 2 | 0.004 | 0.002 | 2.376 | 0.029 | 0.039 |
| **Diet x age** | 2 | 0.003 | 0.002 | 1.950 | 0.023 | 0.064 |
| **Diet x sex** | 1 | 0.001 | 0.001 | 1.047 | 0.006 | 0.326 |
| **Age x sex** | 2 | 0.002 | 0.001 | 1.345 | 0.016 | 0.215 |
| **Diet** | 1 | 0.002 | 0.002 | 2.342 | 0.014 | 0.075 |
| **Age** | 2 | 0.065 | 0.032 | 40.297 | 0.484 | 0.001 |
| **Sex** | 1 | 0.001 | 0.001 | 1.341 | 0.008 | 0.215 |
| **Residuals** | 70 | 0.056 | 0.001 |  | 0.420 |  |
| **Total** | 81 | 0.134 |  |  | 1.000 |  |

Df, degree of freedom; F. Model, F-tests; MeanSqs= SumsOfSqs/df, mean squares; SumsOfSqs, sum of squares. Significance was defined as $P < 0.05$ , by permutational multivariate analysis of variance (PERMANOVA).

A total of 19 families were identified by 16S *rRNA* sequencing. *Lachnospiraceae, Ruminococcaceae* and an unclassified family in the order *Clostridiales* (phylum *Firmicutes),* were the three most abundant families affected by diet, age or sex (Fig.1; Table S2). Exosome-defined diets altered the microbial communities at the family level, whereas some effects depended on diet *x* age and diet *x* age *x* sex interactions. For example, *Anaeroplasmataceae* (phylum of *Tenericutes*; P=0.008), decreased significantly in mice fed ERD compared to mice fed ERS. At age 15 weeks, *Bifidobacteriaceae* (phylum of *Actinobacteria*) and three families (*Lachnospiraceae*, an unclassified family from the order of *Clostridiales* and a family of *Dehalobacteriaceae*) within the phylum of *Firmicutes*, were significantly more abundant in mice fed the ERS diet than mice fed the ERD diet. The family of *Bifidobacteriaceae* was significantly less abundant in mice fed ERS compared with mice fed ERD at age 47 weeks. For diet *x* age *x* sex interactions, *Coriobacteriaceae* (phylum of *Actinobacteria*) and four families (*Ruminococcaceae, Lachnospiraceae, Lactobacillaceae* and *Mogibacteriaceae*) within the phylum of *Firmicutes* were significantly more abundant in male mice fed ERS compared with males fed ERD at age 7 weeks (*P*<0.05). In contrast, the abundance of an unclassified family in the order *Clostridiales* (phylum of *Firmicutes*) was significantly less abundant in female mice fed the ERS diet compared with ERD at age 7 weeks (*P*=0.010). No significant differences were detected at the family level in females and males at age 15 weeks. At age 47 weeks, females fed the ERD diet harbored significantly more *Bifidobacteriaceae* (phylum of *Actinobacteria*; *P*=0.005) and less *Lactobacillaceae* (phylum of *Firmicutes*; *P*=0.018) than ERS females age 47 weeks. Males fed ERD harbored significantly more *Mogibacteriaceae* (phylum of *Firmicutes*; *P*=0.033) and an unclassified family in the order *Clostridiales* (phylum of *Firmicutes*; *P*=0.002) compared with ERS males age 47 weeks.

**FIG 1.**
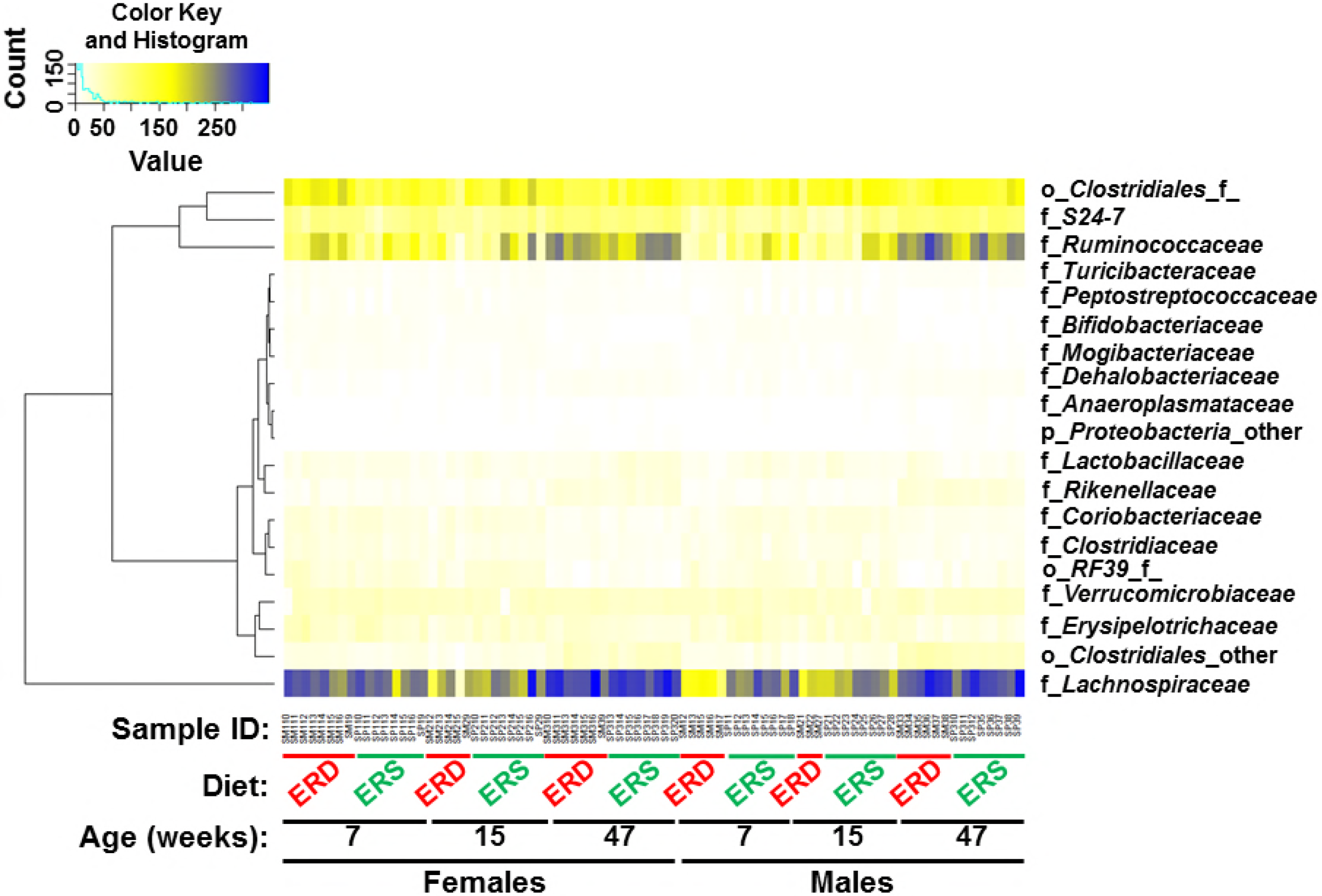
Microbial families (heat map) in the cecum of mice fed exosome RNA-sufficient (ERS) or exosome RNA-depleted (ERD) diets at ages 7, 15 and 47 weeks. f, family; o, order; p, phylum.

At the level of operational taxonomic units (OTUs), the consumption of milk exosome-defined diets had a strong effect on microbial communities in the mouse cecum (Fig. 2). Two OTUs from the family of *Lachnospiraceae* (phylum of *Firmicutes*) were significantly more abundant in mice fed ERS compared with ERD-fed mice at age 7 weeks (*P*=0.009 and *P*=0.044, respectively). At age 15 weeks, OTUs from the families *Verrucomicrobiaceae* (phylum of *Verrucomicrobia*) and *S24-7* (phylum of *Bacteroidetes*) increased significantly in abundance and became the predominant OTUs in mice fed the ERD diet compared to ERS diet (*P*=0.016 and *P*=0.021, respectively). The diversity of microbial OTUs continued to increase and peaked at age 47 weeks and these OTUs are from the same families shown in Fig. 1 in both diet groups. OTUs are detailed in Table S3.

**FIG 2.**
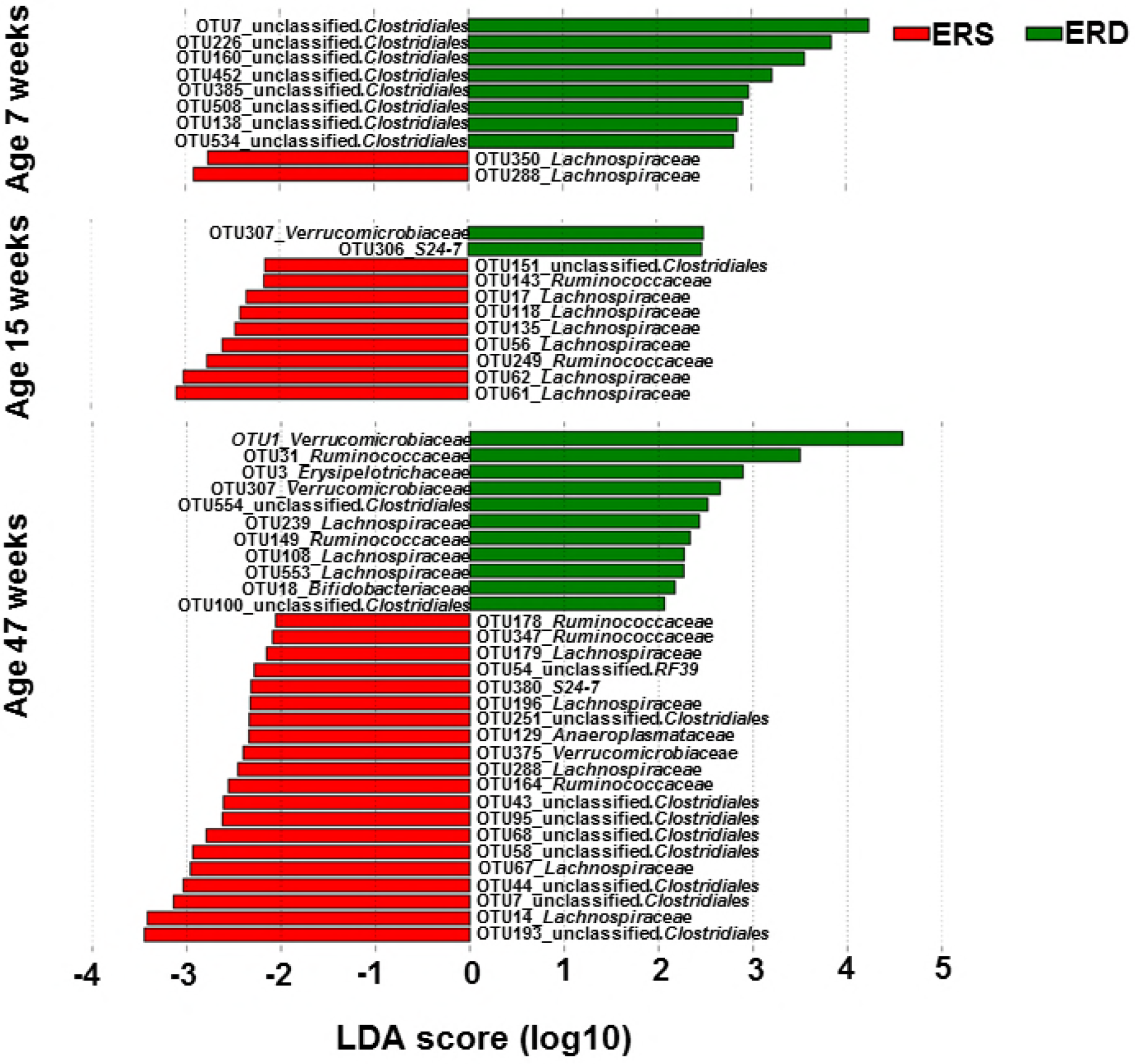
Microbial operational taxonomic units (OTUs) in the cecum of mice fed ERS or ERD diets at ages 7, 15 and 47 weeks (linear discriminant analysis, LDA). Negative (red) and positive (green) LDA scores indicate increased abundance in mice fed ERS and ERD diets, respectively. Effects of diet are statistically significant for LDA scores greater than 2 (*P*<0.05).

### Correlation between microbial communities and hepatic transcriptome

Sixty-nine genes were differentially expressed by at least 2-fold in the livers of female mice fed ERS compared to ERD females at age 15 weeks (*P*<0.01; Table S4). Changes in the hepatic transcriptome correlated with changes in microbial communities in the cecum. The strong correlation between OTUs from the family of *Lachnospiraceae* (phylum of *Firmicutes*) with hepatic transcripts is particularly noteworthy (Fig. 3). For example, five OTUs within the family of *Lachnospiraceae* correlated with the hepatic expression of differentially expressed genes in female mice fed milk exosome-defined diets. Correlations were also observed for OTUs from *Erysipelotrichaceae, Lactobacillaceae*, an unclassified family in the order of *RF39* and four OTUs from an unclassified family in the order *Clostridiales*. All these families, except that from the order of *RF39*, are from the phylum of *Firmicutes.* A full record of correlations is available in Table S5.

**FIG 3.**
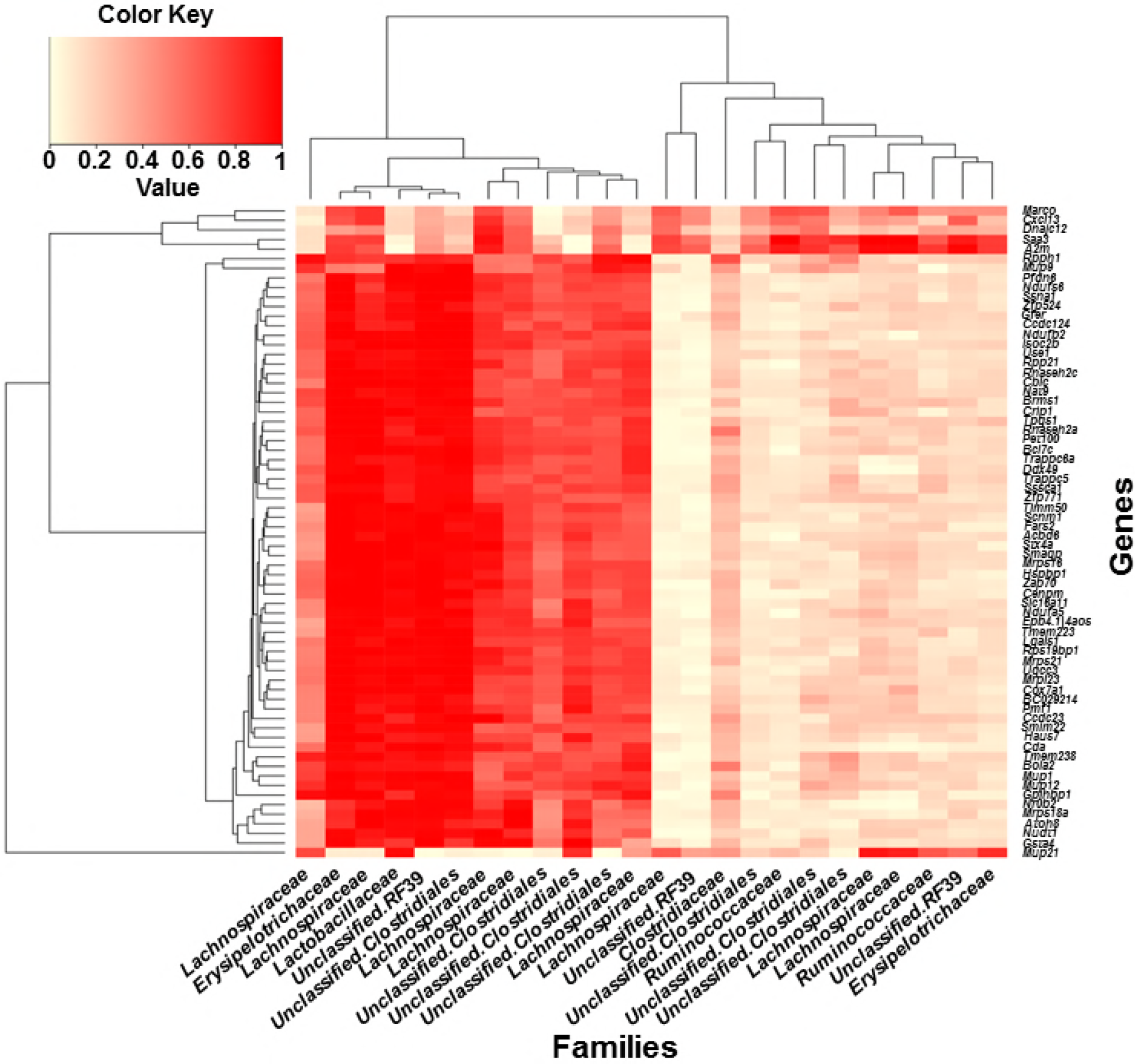
Correlation between changes in OTUs with changes in the hepatic transcriptome in female mice, age 15 weeks fed ERS or ERD (*P*<0.05).TABLE 1 PERMANOVA analysis of the effect of diet, age and sex on microbial communities in the cecum of mice.

## DISCUSSION

This paper provides novel insights in the following areas of research. First, this paper provides strong evidence that dietary exosomes and their RNA cargos, at least those in milk, might be responsible for some of the effects of diet on the gut microbiome (26, 27). To the best of our knowledge the only evidence that bovine milk compounds alter the microbiome comes from a study reporting that consumption of casein protein alters microbial communities in the gut of rats (28). Second, this paper adds to the existing body of evidence that dietary RNAs elicit changes in gene expression across kingdoms, in this case animals and bacteria. This concept is based on a report suggesting that MIR-168a in rice is bioavailable and binds to the mRNA coding for low-density lipoprotein receptor adaptor protein 1, thereby lowering mRNA expression in the liver of mice (29). Third, this paper suggests that gut microorganisms might act as transmitters or amplifiers of dietary exosome signals. Following early reports that microRNA cargos in dietary exosomes might be bioavailable, it was proposed that dietary microRNAs achieve tissue concentrations that are too low to elicit biological effects and that the bioavailability of milk exosomes might be low (15, 16, 30, 31). Here we provide evidence that exosome-defined diets alter microbial communities in the murine cecum, and effects are particularly strong if studied at the OTU level. Importantly, this paper suggests that milk exosomes that escape absorption by mucosal cells, may still elicit major biological effects, facilitated by the gut microbiome. It is now widely accepted that prokaryotic and eukaryotic microbes communicate with their environment through exosome-like vesicles (19–21). It is reasonable to speculate that changes in microbial communities are paralleled by changes in the production of microbial metabolites, which may transmit and amplify milk exosome signals (32). Fourth, this paper provides preliminary evidence that milk exosome-dependent changes in microbial communities can explain changes observed in the hepatic transcriptome in mice. We acknowledge that the scientific basis for this theory is rather small, because this aspect of investigation is based on only one sex (females) in one age group (15 weeks). This being said, we are in the process of integrating exosome-dependent changes in the gut microbiome with changes in the transcriptomes, proteomes and metabolomes in both sexes in various tissues in mice and link the data with changes in murine phenotypes.

The microbial communities altered by milk exosome-defined diets are related to pathological and physiological conditions, as evidenced by the following examples. A loss of *Lachnospiraceae* and *Ruminococcaceae* and a gain in *Enterobacteriaceae* in the ileal mucosa have been implicated in inflammatory bowel disease (33–35). The ratio of *Firmicutes* and *Bacteroidetes* is greater in obese compared with lean subjects (26). Dysbiosis in the gut microbiome may cause liver disease due to microbial metabolites altering the metabolism in hepatic cells via innate immune receptors (36, 37), e.g., a decreased abundance of *Ruminococcaceae* and *Escherichia* has been linked with non-alcoholic fatty liver disease (38, 39). Inflammatory bowel disease, obesity and non-alcoholic fatty liver disease are major health concerns in the United States (40, 41).

Most likely, gut microbiota are not the only amplifiers of exosome and RNA cargo signals. It has been proposed that microRNAs elicit biological effects through binding to Toll-like receptors or by surface antigen-mediated delivery of exosomes to immune cells to create an exosome-rich microenvironment (42, 43); mere exosome-cell surface interactions might also cell signaling pathways (43).

We have recently completed a one-year mouse feeding trial to assess phenotypes associated with feeding milk exosome-defined diets to mice. Some interesting phenotypes emerged (unpublished observations). It will be interesting to test which of these phenotypes depend on the presence of gut microbiota by comparing germ-free and conventional mice or conducting fecal transplant studies.

## MATERIALS AND METHODS

### Mouse diets

Exosome and RNA-depleted (ERD) and exosome and RNA-sufficient (ERS) diets are based on the AIN-93G formulation (6, 44). Lyophilized milk powder (and soy protein) substituted for milk casein in the AIN-93G formulation to eliminate dairy exosomes present in the AIN-93G formulation. The milk added to the diets provided the equivalent of 0.5 L milk consumed by a human adult per day, adjusted by body wright in mice. The milk used to prepare the powder for the ERD diet was ultrasonicated for 1.5 h and incubated for 1 h at 37°C prior to lyophilization; the milk used to prepare the powder for the ERS was not ultrasonicated. Ultrasonication causes a transient disruption of exosome membranes, which led to a >98% depletion of microRNA cargos in exosomes, 20% decreased in exosome count (9.1×10^12^±7.1×10^11^ exosomes/mL in ERS milk vs. 7.3×10^12^±3.5×10^11^ exosomes/mL in ERD milk) and a >60% decrease in intestinal exosome transport rates ((6), S. Sukreet and J. Zempleni, unpublished). Diet ingredients other than milk were not ultrasonicated, i.e., nutrients other than exosomes and their RNA cargos were the same in ERD and ERS diets. Diets were dried and fed in pelleted form.

### Mouse feeding studies

C57BL/6 mice were obtained from Jackson Labs. (stock number 000664) at age three weeks when dietary treatment was initiated ERD versus ERS. Mice were housed in groups of four mice per cage, separated by sex. Both males and females were studied. True randomization of group assignment was achieved by labeling mice with numbers and randomly assigning numbers to groups. At timed intervals (ages 7, 15 and 47 weeks), mice were sampled from different cages to avoid cage effects, and euthanized for sample collection (N=8 for each sex and age). Cecum content was collected, flash frozen in liquid nitrogen and stored at −80°C. Livers were collected at age 15 weeks and flash frozen in liquid nitrogen and stored at - 80°C. The study was approved by the Institutional Animal Care and Use Committee at the University of Nebraska-Lincoln (protocol 1229).

### Analysis of microbial communities

Cecum content was extracted and DNA was purified using the PowerSoil DNA Isolation Kit (Mo Bio Laboratories Inc., Carlsbad, CA, USA) following the manufacturer’s instructions. DNA purity and integrity were confirmed by using the 260-to-280 nm ratio (Nanodrop ND-1000, Nanodrop Technologies, Wilmington, DE, USA) and agarose gel (0.8%) electrophoresis. The V4 region in the *16S rRNA* gene was amplified and sequenced as described previously (45). The sequencing reads were quality filtered and analyzed as described previously (46). Briefly, contigs were generated from paired-end reads and were screened using MOTHUR v.1.38.1 (46) to exclude low quality sequences and reads containing ambiguous bases or homopolymers longer than 8 bp. Additionally, the resulting reads were trimmed to only retain reads between 245 base pairs (bp) and 275 bp. The UPARSE pipeline (USEARCH v7.0.1090) (47) was then used to cluster quality-filtered sequences into operational taxonomic units (OTUs) at 97% identity, after removal of chimeras using UCHIME (48). ChimeraSlayer gold.fa was used as the reference database for chimera detection. Sequence alignment was performed using the SILVA v123 reference and was used to build a phylogenetic tree using Clearcut (49). Taxonomy assignment (Greengenes database: gg_13_8_otus) (50) was performed using QIIME v.1.9.1 (51). We removed 10 samples that had sequencing read counts lower than 4000. Eighty-two samples with an average read count of 40,468 reads and a range of 4,112 – 144,788 reads were used for downstream analysis. Alpha diversity metrics were used to evaluate richness (Chao1), diversity (Shannon-Weiner index), and coverage (Good’s coverage) (51). Rarefaction curves were constructed using Chao1 values. A core measureable microbiome was identified based on factors diet, sex, and age. The core measureable microbiome was defined as the group of OTUs that are present in at least 80% of the samples within each factor. Differences in bacterial communities were assessed using permutational multivariate analysis of variance (PERMANOVA) utilizing the weighted UniFrac distance matrix. Additionally, the weighted UniFrac distance matrix was used for principal coordinate analysis (PCoA). The Linear Discriminant Analysis of Effect Size (LefSe) algorithm with default parameters was used to identify OTUs that were differentially abundant in the ERS and ERD feeding groups at different ages (52). Sequence data were deposited in the NCBI-BioProject database under accession no. PRJNA413623.

### Analysis of the hepatic transcriptome

Livers from six female mice, age 15 weeks, were used for RNA sequencing studies (N=3 from each feeding group). Briefly, RNA was extracted using the mRNA Seq Sample Prep Kit (Illumina) and shipped on dry ice to the genomics center of University of Minneapolis, MN for sequencing analysis. RNA quality was assessed using an Agilent Bioanalyzer and absorbance at 260 and 280 nm. The RNA Integrity Number and the 260-to-280 nm ratio was greater than 7 and 1.8, respectively, for all samples. Libraries were generated using TruSeq Stranded Total RNA Library Prep Kit and sequenced using the Illumina HiSeq 2500 platform and a paired ends protocol generating reads with a length of 125 bp. Data quality control was performed using FastQC (53). After removing adaptors and reads containing ambiguous bases or having average quality score less than 30, sequencing reads were aligned to the mouse reference genome [GRCm38, mm10] using Tophat (54). Cufflinks (55) was applied to identify the transcripts and quantify their expression in units of reads per kilobase of exon model per million (RPKM). Cuffdiff was applied to identify the differentially expressed transcripts between two feeding groups and only those with equal or more than 2-fold change were considered for the downstream analysis (56). KEGG pathways were identified by using clusterProfiler (57). Raw sequencing data were deposited in the NCBI-BioProject database under accession no. PRJNA400248.

### Correlation analysis of microbial communities and hepatic transcriptome

Correlations between the significantly differentially abundant OTUs and the top 69 significantly differentially expressed genes were calculated using the Pearson product-moment correlation and bootstrapping with 1000 permutations to calculate the p-values of the correlation scores (QIIME v.1.9.1). To visualize the microbial features that correlated with gene expression, a heatmap was generated using R software package 3.3.3 (The R Foundation). Bray-Curtis dissimilarity was used to calculate the distance for rows and columns, and average linkage hierarchical clustering was used to generate dendrograms.

### Statistical analysis

Kruskal-Wallis sum-rank test was used to identify the significant differences in abundance between groups at *P*<0.05. The Wilcoxon rank-sum test was used for pairwise comparisons at adjusted *P*<0.05, with the Benjamini and Hochberg correction (58). Differences were considered statistically significant if *P*<0.05.

### Data availability

Sequencing data have been deposited in the NCBI-BioProject database under accession no. PRJNA413623 (microbiome) and PRJNA400248 (transcriptome), respectively.

## SUPPLEMENTAL MATERIAL

**TABLE S1** Kruskal-Wallis rank sum test for the effects of age, sex and diet effect on microbial phyla

**TABLE S2** Kruskal-Wallis rank sum test for the effects of age, sex and diet effect on the 19 most abundant families

**TABLE S3** LefSe (linear discriminant analysis coupled with effect size measurements) test for the effects of age and diet effect on microbial families

**TABLE S4** Differentially expressed genes in livers from female C57BL/6 mice fed milk exosome defined diets

**TABLE S5** Correlation between OTUs with hepatic transcripts in female C57BL/6 mice fed milk exosome defined diets

## ACKNOWLEDGMENTS

We acknowledge the support of the Biomedical and Obesity Research Core in the Nebraska Center for the Prevention of Obesity Diseases through Dietary Molecules (NIH 1P20GM104320) and the Holland Computing Center.

This work was supported by the National Institute of Food and Agriculture (NIFA) 201567017-23181 and 2016-67001-25301, the National Institutes of Health 1P20GM104320, the Gerber Foundation, the Egg Nutrition Center/United States Department of Agriculture, the University of the Nebraska President’s Office, the University of Nebraska Agricultural Research Division Hatch Act, the United States Department of Agriculture multistate group W3002 (all to JZ), and the University of Nebraska Agricultural Research Division, Hatch Act, an ARS/USMARC collaborative funding program, and the United States Department of Agriculture multistate group W2010 (all to SF).

FZ made substantial contributions to the acquisition, analysis and interpretation of data and wrote the first version of the manuscript. HP made substantial contributions to the acquisition of data and was involved in revising the manuscript. JS made substantial contributions to the analysis and interpretation of data and was involved in revising the manuscript. MS made substantial contributions to the acquisition of samples and data. JC made substantial contributions to the analysis and interpretation of data and was involved in writing and revising the manuscript. SF made substantial contributions to the analysis and interpretation of data and was involved in writing and revising the manuscript. JZ conceived the idea for this study, made substantial contributions to the interpretation of data and wrote the final version of the manuscript. All authors have approved the final version of the manuscript and will take public responsibility for appropriate portions of the content. All authors have agreed to be accountable for all aspects of the work in ensuring that questions related to the accuracy or integrity of any part of the work are appropriately investigated and resolved.

JZ is a consultant for PureTech Health, Inc. in Boston, MA. SF is a co-owner of NuGUT LLC. The other authors declare no conflict of interest.

## REFERENCES

1. Yanez-Mo M, Siljander PR, Andreu Z, Zavec AB, Borras FE, Buzas EI, Buzas K, Casal E, Cappello F, Carvalho J, Colas E, Cordeiro-da Silva A, Fais S, Falcon-Perez JM, Ghobrial IM, Giebel B, Gimona M, Graner M, Gursel I, Gursel M, Heegaard NH, Hendrix A, Kierulf P, Kokubun K, Kosanovic M, Kralj-Iglic V, Kramer-Albers EM, Laitinen S, Lasser C,Lener T, Ligeti E, Line A, Lipps G, Llorente A, Lotvall J, Mancek-Keber M, Marcilla A, Mittelbrunn M, Nazarenko I, Nolte-’t Hoen EN, Nyman TA, O’Driscoll L, Olivan M, Oliveira C, Pallinger E, Del Portillo HA, Reventos J, Rigau M, Rohde E, Sammar M, et al. 2015. Biological properties of extracellular vesicles and their physiological functions. J Extracell Vesicles 4:27066.

2. Abels ER, Breakefield XO. 2016. Introduction to Extracellular Vesicles: Biogenesis, RNA Cargo Selection, Content, Release, and Uptake. Cell Mol Neurobiol 36:301.

3. Vilella F, Moreno-Moya JM, Balaguer N, Grasso A, Herrero M, Martinez S, Marcilla A, Simon C. 2015. Hsa-miR-30d, secreted by the human endometrium, is taken up by the preimplantation embryo and might modify its transcriptome. Development 142:3210–3221.

4. Wolf T, Baier SR, Zempleni J. 2015. The intestinal transport of bovine milk exosomes is mediated by endocytosis in human colon carcinoma caco-2 cells and rat small intestinal IEC-6 cells. J Nutr 145:2201–2206.

5. Kusuma Jati R, Manca S, Friemel T, Sukreet S, Nguyen C, Zempleni J. 2016. Human vascular endothelial cells transport foreign exosomes from cow’s milk by endocytosis.. Am J Physiol Cell Physiol 310:C800–C807.

6. Baier SR, Nguyen C, Xie F, Wood JR, Zempleni J. 2014. MicroRNAs are absorbed in biologically meaningful amounts from nutritionally relevant doses of cow’s milk and affectgene expression in peripheral blood mononuclear cells, HEK-293 kidney cell cultures, and mouse livers. J Nutr 144:1495–1500.

7. Izumi H, Tsuda M, Sato Y, Kosaka N, Ochiya T, Iwamoto H, Namba K, Takeda Y. 2015. Bovine milk exosomes contain microRNA and mRNA and are taken up by human macrophages. J Dairy Sci 98:2920–2933.

8. Imai T, Takahashi Y, Nishikawa M, Kato K, Morishita M, Yamashita T, Matsumoto A, Charoenviriyakul C, Takakura Y. 2015. Macrophage-dependent clearance of systemically administered B16BL6-derived exosomes from the blood circulation in mice. J Extracell Vesicles 4:26238.

9. Shu J, Chiang K, Zempleni J, Cui J. 2015. Computational characterization of exogenous microRNAs that can be transferred into human circulation. PLoS One 10:e0140587.

10. Wang L, Sadri M, Giraud D, Zempleni J. 2018. RNase H2-Dependent Polymerase Chain Reaction and Elimination of Confounders in Sample Collection, Storage, and Analysis Strengthen Evidence That microRNAs in Bovine Milk Are Bioavailable in Humans. J Nutr 148:153–159.

11. Zhou F, Paz AH, Sadri M, Fernando CS, Zempleni J. 2017. A diet defined by its content of bovine milk exosomes alters the composition of the intestinal microbiome in C57BL/6 mice. FASEB J 31:965.24 [peer-reviewed meeting abstract].

12. Sadri M, Xie F, Wood J, Zempleni J. 2016. Dietary depletion of cow’s milk microRNAs impairs fecundity in mice. FASEB J 30:673.5. [peer-reviewed meeting abstract].

13. Mutai E, Zhou F, Zempleni J. 2017. Depletion of dietary bovine milk exosomes impairs sensorimotor gating and spatial learning in C57BL/6 mice. FASEB J 31:150.4. [peer-reviewed meeting abstract].

14. Aguilar-Lozano A, Baier SR, Adamec J, Sadri M, Giraud D, Zempleni J. 2016. Depletion of dietary microRNAs from cow’s milk causes an increase of purine metabolites in human body fluids and mouse livers. FASEB J 30 (supplement 1): 127.1. [peer-reviewed meeting abstract].

15. Laubier J, Castille J, Le Guillou S, Le Provost F. 2015. No effect of an elevated miR-30b level in mouse milk on its level in pup tissues. RNA Biol 12:26–29.

16. Title AC, Denzler R, Stoffel M. 2015. Uptake and function studies of maternal milk-derived microRNAs. J Biol Chem 290:23680–23691.

17. Zempleni J, Aguilar-Lozano A, Sadri M, Sukreet S, Manca S, Wu D, Zhou F, Mutai E. 2017. Biological activities of extracellular vesicles and their cargos from bovine and human milk in humans and implications for infants. J Nutr 147:3–10.

18. Zempleni J. 2017. Milk exosomes: beyond dietary microRNAs. Genes Nutr 12:12.

19. Wolf JM, Casadevall A. 2014. Challenges posed by extracellular vesicles from eukaryotic microbes. Curr Opin Microbiol 22C:73–78.

20. Rivera J, Cordero RJ, Nakouzi AS, Frases S, Nicola A, Casadevall A. 2010. Bacillus anthracis produces membrane-derived vesicles containing biologically active toxins. Proc Natl Acad Sci USA 107:19002–19007.

21. Lee J, Lee EY, Kim SH, Kim DK, Park KS, Kim KP, Kim YK, Roh TY, Gho YS. 2013. Staphylococcus aureus extracellular vesicles carry biologically active beta-lactamase. Antimicrob Agents Chemother 57:2589–2595.

22. Meckes DG Jr., 2015. Exosomal communication goes viral. J Virol 89:5200–5203.

23. Wang K, Li H, Yuan Y, Etheridge A, Zhou Y, Huang D, Wilmes P, Galas D. 2012. The complex exogenous RNA spectra in human plasma: an interface with human gut biota? PLoS One 7:e51009.

24. Munagala R, Aqil F, Jeyabalan J, Gupta RC. 2016. Bovine milk-derived exosomes for drug delivery. Cancer Lett 371:48–61.

25. Manca S, Giraud D, Zempleni J. 2017. The bioavailability and distribution of bovine milk exosomes is distinct from that of their cargos in mice. FASEB J 31:148.2 [peer-reviewed meeting abstract].

26. Ley RE, Turnbaugh PJ, Klein S, Gordon JI. 2006. Microbial ecology: human gut microbes associated with obesity. Nature 444:1022–1023.

27. Wu GD, Chen J, Hoffmann C, Bittinger K, Chen YY, Keilbaugh SA, Bewtra M, Knights D, Walters WA, Knight R, Sinha R, Gilroy E, Gupta K, Baldassano R, Nessel L, Li H, Bushman FD, Lewis JD. 2011. Linking long-term dietary patterns with gut microbial enterotypes. Science 334:105–108.

28. Zhu Y, Lin X, Zhao F, Shi X, Li H, Li Y, Zhu W, Xu X, Li C, Zhou G. 2015. Meat, dairy and plant proteins alter bacterial composition of rat gut bacteria. Sci Rep 5:15220.

29. Zhang L, Hou D, Chen X, Li D, Zhu L, Zhang Y, Li J, Bian Z, Liang X, Cai X, Yin Y, Wang C, Zhang T, Zhu D, Zhang D, Xu J, Chen Q, Ba Y, Liu J, Wang Q, Chen J, Wang J, Wang M, Zhang Q, Zhang J, Zen K, Zhang CY. 2012. Exogenous plant MIR168a specifically targets mammalian LDLRAP1: evidence of cross-kingdom regulation by microRNA. Cell Res 22:107–126.

30. Snow JW, Hale AE, Isaacs SK, Baggish AL, Chan SY. 2013. Ineffective delivery of diet-derived microRNAs to recipient animal organisms. RNA Biol 10:1107–1116.

31. Dickinson B, Zhang Y, Petrick JS, Heck G, Ivashuta S, Marshall WS. 2013. Lack of detectable oral bioavailability of plant microRNAs after feeding in mice. Nat Biotechnol 31:965–967.

32. Wikoff WR, Anfora AT, Liu J, Schultz PG, Lesley SA, Peters EC, Siuzdak G. 2009. Metabolomics analysis reveals large effects of gut microflora on mammalian blood metabolites. Proc Natl Acad Sci USA 106:3698–3703.

33. Frank DN, St Amand AL, Feldman RA, Boedeker EC, Harpaz N, Pace NR. 2007. Molecular-phylogenetic characterization of microbial community imbalances in human inflammatory bowel diseases. Proc Natl Acad Sci USA 104:13780–13785.

34. Fujimoto T, Imaeda H, Takahashi K, Kasumi E, Bamba S, Fujiyama Y, Andoh A. 2013. Decreased abundance of Faecalibacterium prausnitzii in the gut microbiota of Crohn’s disease. J Gastroenterol Hepatol 28:613–619.

35. Baumgart M, Dogan B, Rishniw M, Weitzman G, Bosworth B, Yantiss R, Orsi RH, Wiedmann M, McDonough P, Kim SG, Berg D, Schukken Y, Scherl E, Simpson KW. 2007. Culture independent analysis of ileal mucosa reveals a selective increase in invasive Escherichia coli of novel phylogeny relative to depletion of Clostridiales in Crohn’s disease involving the ileum. ISME J 1:403–418.

36. Marchesi JR, Adams DH, Fava F, Hermes GD, Hirschfield GM, Hold G, Quraishi MN, Kinross J, Smidt H, Tuohy KM, Thomas LV, Zoetendal EG, Hart A. 2016. The gut microbiota and host health: a new clinical frontier. Gut 65:330–339.

37. Adams DH, Eksteen B, Curbishley SM. 2008. Immunology of the gut and liver: a love/hate relationship. Gut 57:838–848.

38. Raman M, Ahmed I, Gillevet PM, Probert CS, Ratcliffe NM, Smith S, Greenwood R, Sikaroodi M, Lam V, Crotty P, Bailey J, Myers RP, Rioux KP. 2013. Fecal microbiome and volatile organic compound metabolome in obese humans with nonalcoholic fatty liver disease. Clin Gastroenterol Hepatol 11:868-875 e861–863.

39. Zhu L, Baker SS, Gill C, Liu W, Alkhouri R, Baker RD, Gill SR. 2013. Characterization of gut microbiomes in nonalcoholic steatohepatitis (NASH) patients: a connection between endogenous alcohol and NASH. Hepatology 57:601–609.

40. Centers for Disease Control and Prevention. 2011. Inflammatory bowel disease, 01/11/2011; 2011. http://www.cdc.gov/ibd/ (accessed: 4/8/2013).

41. Centers for Disease Control and Prevention. 2015. Obesity and Overweight. http://www.cdc.gov/nchs/fastats/obesity-overweight.htm (accessed: 6/1/2017).

42. Fabbri M, Paone A, Calore F, Galli R, Gaudio E, Santhanam R, Lovat F, Fadda P, Mao C, Nuovo GJ, Zanesi N, Crawford M, Ozer GH, Wernicke D, Alder H, Caligiuri MA, Nana-Sinkam P, Perrotti D, Croce CM. 2012. MicroRNAs bind to Toll-like receptors to induce prometastatic inflammatory response. Proc Natl Acad Sci USA 109:E2110–E2116.

43. Bryniarski K, Ptak W, Jayakumar A, Pullmann K, Caplan MJ, Chairoungdua A, Lu J,Adams BD, Sikora E, Nazimek K, Marquez S, Kleinstein SH, Sangwung P, Iwakiri Y, Delgato E, Redegeld F, Blokhuis BR, Wojcikowski J, Daniel AW, Groot Kormelink T, Askenase PW. 2013. Antigen-specific, antibody-coated, exosome-like nanovesicles deliver suppressor T-cell microRNA-150 to effector T cells to inhibit contact sensitivity. J Allergy Clin Immunol 132:170–181.

44. Reeves PG, Nielsen FH, Fahey GC Jr., 1993. AIN-93 purified diets for laboratory rodents: final report of the American Institute of Nutrition ad hoc writing committee on the reformulation of the AIN-76A rodent diet. J Nutr 123:1939–1951.

45. Kozich JJ, Westcott SL, Baxter NT, Highlander SK, Schloss PD. 2013. Development of a dual-index sequencing strategy and curation pipeline for analyzing amplicon sequence data on the MiSeq Illumina sequencing platform. Appl Environ Microbiol 79:5112–5120.

46. Schloss PD, Westcott SL, Ryabin T, Hall JR, Hartmann M, Hollister EB, Lesniewski RA, Oakley BB, Parks DH, Robinson CJ, Sahl JW, Stres B, Thallinger GG, Van Horn DJ,Weber CF. 2009. Introducing mothur: open-source, platform-independent, community-supported software for describing and comparing microbial communities. Appl Environ Microbiol 75:7537–7541.

47. Edgar RC. 2013. UPARSE: highly accurate OTU sequences from microbial amplicon reads. Nat Methods 10:996–998.

48. Edgar RC, Haas BJ, Clemente JC, Quince C, Knight R. 2011. UCHIME improves sensitivity and speed of chimera detection. Bioinformatics 27:2194–2200.

49. Sheneman L, Evans J, Foster JA. 2006. Clearcut: a fast implementation of relaxed neighbor joining. Bioinformatics 22:2823–2824.

50. McDonald D, Price MN, Goodrich J, Nawrocki EP, DeSantis TZ, Probst A, Andersen GL, Knight R, Hugenholtz P. 2012. An improved Greengenes taxonomy with explicit ranks for ecological and evolutionary analyses of bacteria and archaea. ISME J 6:610–618.

51. Caporaso JG, Kuczynski J, Stombaugh J, Bittinger K, Bushman FD, Costello EK, Fierer N, Pena AG, Goodrich JK, Gordon JI, Huttley GA, Kelley ST, Knights D, Koenig JE, Ley RE, Lozupone CA, McDonald D, Muegge BD, Pirrung M, Reeder J, Sevinsky JR, Turnbaugh PJ, Walters WA, Widmann J, Yatsunenko T, Zaneveld J, Knight R. 2010. QIIME allows analysis of high-throughput community sequencing data. Nat Methods 7:335–336.

52. Segata N, Izard J, Waldron L, Gevers D, Miropolsky L, Garrett WS, Huttenhower C. 2011. Metagenomic biomarker discovery and explanation. Genome Biol 12:R60.

53. Babraham B. 2017. FastQC. http://www.bioinformatics.babraham.ac.uk/projects/fastqc/ (accessed: 8/24/2017).

54. Trapnell C, Pachter L, Salzberg SL. 2009. TopHat: discovering splice junctions with RNA-Seq. Bioinformatics 25:1105–1111.

55. Roberts A, Pimentel H, Trapnell C, Pachter L. 2011. Identification of novel transcripts in annotated genomes using RNA-Seq. Bioinformatics 27:2325–2329.

56. Trapnell C, Hendrickson DG, Sauvageau M, Goff L, Rinn JL, Pachter L. 2013. Differential analysis of gene regulation at transcript resolution with RNA-seq. Nat Biotechnol 31:46–53.

57. Yu G, Wang LG, Han Y, He QY. 2012. clusterProfiler: an R package for comparing biological themes among gene clusters. OMICS 16:284–287.

58. Benjamini Y, Hochberg Y. 1995. Controlling the False Discovery Rate – a Practical and Powerful Approach to Multiple Testing. J Roy Stat Soc Ser B Methodol 57:289–300.

